# Biophysical evolution of the receptor binding domains of SARS-CoVs

**DOI:** 10.1101/2023.06.13.544630

**Authors:** Vaibhav Upadhyay, Sudipta Panja, Alexandra Lucas, Casey Patrick, Krishna M.G. Mallela

## Abstract

With hundreds of coronaviruses (CoVs) identified in bats that are capable of infecting humans, it is important to understand how CoVs that affected the human population have evolved. Seven known coronaviruses have infected humans, of which three CoVs caused severe disease with high mortality rates: SARS-CoV emerged in 2002, MERS-CoV in 2012, and SARS-CoV-2 in 2019. Both SARS-CoV and SARS-CoV-2 belong to the same family, follow the same receptor pathway, and use their receptor binding domain (RBD) of spike protein to bind to the ACE2 receptor on the human epithelial cell surface. The sequence of the two RBDs is divergent, especially in the receptor binding motif (RBM) that directly interacts with ACE2. We probed the biophysical differences between the two RBDs in terms of their structure, stability, aggregation, and function. Since RBD is being explored as an antigen in protein subunit vaccines against CoVs, determining these biophysical properties will also aid in developing stable protein subunit vaccines. Our results show that despite RBDs having a similar three-dimensional structure, they differ in their thermodynamic stability. RBD of SARS-CoV-2 is significantly less stable than that of SARS-CoV. Correspondingly, SARS-CoV-2 RBD shows a higher aggregation propensity. Regarding binding to ACE2, less stable SARS-CoV-2 RBD binds with a higher affinity than more stable SARS-CoV RBD. In addition, SARS-CoV-2 RBD is more homogenous in terms of its binding stoichiometry towards ACE2, compared to SARS-CoV RBD. These results indicate that SARS-CoV-2 RBD differs from SARS-CoV RBD in terms of its stability, aggregation, and function, possibly originating from the diverse RBMs. Higher aggregation propensity and decreased stability of SARS-CoV-2 RBD warrants further optimization of protein subunit vaccines that use RBD as an antigen either by inserting stabilizing mutations or formulation screening.

**Statement of Significance:** This study holds significant relevance in the context of the COVID-19 pandemic and the broader understanding of coronaviruses. A comparison of the receptor binding domains (RBDs) of SARS-CoV and SARS-CoV-2 reveals significant differences in their structure, stability, aggregation, and function. Despite divergent sequences, the RBDs share a similar fold and ACE2 receptor binding capability, likely through convergent evolution. These findings are crucial for understanding coronavirus evolution, interactions with human receptors, and the spillover of coronaviruses from animals to humans. The study also has implications for vaccine design strategies for SARS-CoVs, where the RBD is used as an antigen in protein subunit vaccines. By anticipating future outbreaks and enhancing our understanding of zoonotic spillover, this research contributes to safeguarding human health.

## INTRODUCTION

Coronavirus disease 2019 (COVID-19) caused by SARS-CoV-2 emerged in Wuhan, China in December 2019 and spread around the world as a global pandemic affecting more than 180 countries. (1-3) As of June 2023, the pandemic has infected about 690 million people causing 6.9 million deaths making it one of the worst pandemics of all time. Similar outbreaks by coronaviruses, although less severe in terms of the number of cases, occurred in 2002 and 2012. The causative agents for these outbreaks were SARS-CoV and MERS-CoV. These three coronaviruses belong to the Betacoronavirus genus of the family Coronaviridae, which contains single-stranded RNA as its genetic material.(4) It is widely considered that human coronaviruses are derived from the bat reservoir,(5, 6) although the specific origin of SARS-CoV-2 is still being debated.(7, 8) To be able to infect human hosts, the virus needs to acquire features necessary for infecting humans. SARS-CoV-2 and SARS-CoV belong to the sarbecovirus subgenus and utilize their spike protein’s receptor binding domain (RBD) to interact with the ACE2 receptor on the human epithelial cells and gain cellular entry.(9-13) MERS-CoV, on the other hand, uses its spike RBD to interact with dipeptidyl peptidase 4 (DPP4) to enter human cells.(14, 15)

The mutations acquired by RBD during the evolution of SARS-CoV-2 from its precursors are important in dictating its interaction with ACE2 and determining its potential to infect humans. The host range of the virus is largely determined by these mutations.(16) SARS-CoV acquired RBD mutations that enabled it to interact with ACE2, probably through a different evolutionary trajectory.(17) The evolution of similar ACE2 binding function in SARS-CoV and SARS-CoV-2 RBDs points towards convergent evolution and raises the possibility of future outbreaks of viruses with similar acquisition of traits with human infection potential. It is thus necessary to understand the sequence features of RBD that allow it to interact with ACE2 and understand the structural basis of ACE2-RBD interactions in general. RBD is also important from the viewpoint of immune responses as most of the neutralizing antibody response of the human host is generated against the RBD, and the antibodies from patients recovered from COVID-19 have been developed as therapeutics.(18) Many of the protein subunit vaccine candidates use RBD as the source of antigen.(19) RBD-based nanoparticles have also been shown to bind more efficiently to neutralizing antibodies and activate complement proteins.(20) Antigen stability is an important factor that determines the success of the protein subunit vaccines,(21, 22), and hence studying the molecular properties of RBDs is important in developing stable vaccines against SARS-CoVs.

In this study, we analyzed the biophysical properties of the RBDs of SARS-CoV and SARS-CoV-2 (hereby referred to as RBD1 and RBD2 respectively) and compared them with respect to their sequence, structure, stability, aggregation, and function. Understanding the differences between the two RBDs will also help in coming up with better countermeasures against COVID-19 and any future coronavirus outbreaks, as many of the known countermeasures including vaccines and antibody therapies are based on RBD.(23-28) This approach of understanding the structural and functional characteristics of RBDs becomes important in understanding the emerging new SARS-CoV-2 variants with increased infectivity, many of which have mutations in the RBD.(29)

## MATERIALS AND METHODS

### Protein expression and purification

The genes encoding the N-terminal peptidase domain (19S-615D) of ACE2 (UniProtKB ID-Q9BYF1, NCBI Gene ID-59272) and receptor binding domain of SARS-CoV (333T-541F; UniProtKB ID-P0DTC2, NCBI Gene ID-1489668) and SARS-CoV-2 (306R-527F; UniProtKB ID-P59594, NCBI Gene ID-43740568) were synthesized by Twist Biosciences. Human immunoglobulin heavy chain signal sequence and His-SUMOstar tag were attached towards the 5’ end of the gene and cloned into pcDNA3.4 TOPO vector. The vector was transfected using PEI (40 kDa, linear polymer) into Expi293F cells from Thermo Fisher Scientific for transient expression. The cells were harvested 6 days after transfection and the supernatant was loaded on the Ni-NTA column for affinity purification. The purified tagged protein was digested with SUMOstar protease to remove the His-SUMOstar tag and passed again through the Ni-NTA column to obtain untagged protein in the flow-through and wash fractions. The proteins were dialyzed extensively in pH 7.0 buffer containing 50 mM sodium Phosphate and 20 mM NaCl before use.

### Circular Dichroism (CD) and fluorescence spectroscopy

All CD experiments were performed on the Applied Photophysics Chirascan Plus spectrophotometer. For CD spectra, 20 μM of protein was taken in a 0.5 mm pathlength cuvette, and spectra were recorded from 260 - 185 nm with a data interval of 1 nm and averaging time of 2 s at every wavelength. All Fluorescence experiments were performed on the PTI QuantaMaster fluorimeter. For fluorescence spectra, a 2 μM protein sample was excited at 295 nm and the fluorescence emission spectrum was recorded from 310 - 450 nm with a data interval of 1 nm and averaging time of 1 s/nm.

### Thermal denaturation melts

The thermal denaturation melts were recorded using CD spectroscopy. Spectra were obtained at a protein concentration of 10 μM in a 0.5 mm pathlength cuvette. CD spectra were recorded from 190 - 260 nm with a data interval of 1 nm and an averaging time of 2 s/nm. A step gradient ranging from 20°C to 95°C with a 5°C interval and equilibration time of 5 min at each temperature was employed. The thermal denaturation melts at 222 nm were recorded at a protein concentration of 20 μM in a 0.5 mm pathlength cuvette. A continuous gradient ranging from 20°C to 90°C at a ramp rate of 1°C/min was employed. The data were acquired for every 1°C increment after averaging for 2 s, and were fitted to a two-state unfolding equation,

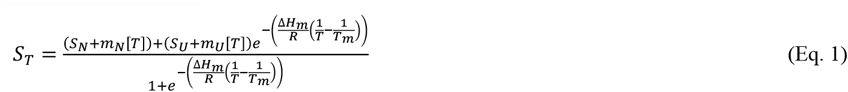

where S_T_ is the measured signal as a function of temperature [T], S_N_ and S_U_ are the signals corresponding to the native and unfolded baselines, m_N_ and m_U_ are the slopes of linear dependence of S_N_ and S_U_, ΔH_m_ is the enthalpy change at T_m_, R is the universal gas constant, and T is the absolute temperature in Kelvin.

### Urea denaturation melts

Urea denaturation melts were recorded using CD and fluorescence spectroscopy. The denaturation melts were obtained by mixing native and denatured protein samples using a Microlab 600 Hamilton syringe titrator at a protein concentration of 2 μM to achieve urea concentrations ranging from 0 to 8.6 M. The incubation time after mixing the two solutions before data acquisition was 5 min. For fluorescence, samples were excited at 280 nm and the emission intensity at 320 nm was used to plot the denaturation curves. For CD, the signal at 222 nm was plotted with respect to the urea concentration to obtain denaturation curves. The denaturation curves were fitted to the following equation,

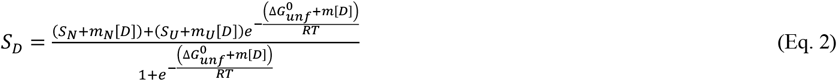

where S_D_ is the measured signal as a function of denaturant concentration [D], S_N_ and S_U_ are the signals corresponding to the native and unfolded baselines, m_N_ and m_U_ are the slopes of linear dependence of S_N_ and S_U_, ΔG^0^_unf_ is the Gibbs free energy of unfolding from native to unfolded state in the absence of denaturant, m is the slope of linear variation of Gibbs free energy with denaturant concentration, R is the universal gas constant, and T is the temperature in Kelvin.

### Protein aggregation kinetics

Aggregation kinetics of RBDs were recorded using a PTI fluorescence spectrometer with excitation and emission monochromators set at 350 nm. The protein at a final concentration of 5 μM was added to the buffer pre-heated to 60°C.

### Size determination of protein aggregates

The sizes of the microparticles were determined using FlowCAM 8000 (Fluid-Imaging Technologies, Scarborough, ME) flow microscopy instrument. 200 μL sample volume was injected per run at a flow rate of 0.15 mL/min. Particle count measurements were carried out without centrifugation or filtration. The lowest detection limit of the counting particles was 1 μm and particle distributions were analyzed between 1 and 25 μm window (equivalent spherical diameter). Four successive measurements were performed to determine the mean value of size distribution.

### Size Exclusion Chromatography-Multi Angle Light Scattering (SEC-MALS)

SEC experiments were performed on an Agilent 1100 series chromatographic system equipped with a UV detector. SEC data analysis was done on Chemstation data analysis software. 100 μl of the protein samples in the concentration range of 10-80 μM were loaded onto the TSKgel G3000SWXL column (Tosoh Biosciences) pre-equilibrated with 50 mM Sodium phosphate buffer, 20 mM NaCl, pH 7.0. The flow rate was kept at 0.5 ml/min and the elution of proteins was followed by monitoring absorbance at 280 nm. For SEC-MALS experiments, the proteins eluted from the column were allowed to pass through two additional detectors, a multi-angle light scattering detector and a refractive index detector (Viscotek). For molecular weight estimation of RBDs, a protein concentration of 100 μM was used. For ACE2-RBD protein complex analysis, ACE2 concentration was 24 μM and RBD concentration was 48 μM. OmniSEC-Bio 5.10 software was used to analyze the SEC-MALS data and molecular weight calculations.

### Analytical ultracentrifugation

All sedimentation velocity analytical ultracentrifugation (SV-AUC) experiments were performed using Beckman XL-A ultracentrifuge. 400 μl of the sample and 410 μl of reference buffer (50 mM Phosphate buffer, pH 7.0, and 20 mM NaCl) were transferred to the double-sectored 12 mm thick centerpieces of assembled AUC cells. Absorbance at 280 nm was selected to record the sedimentation scans. The SV-AUC profiles were obtained for ACE2 (5 μM), RBD1 and RBD2 (10 μM each) at a speed of 50,000 rpm. The SV-AUC profiles for the ACE2-RBD complexes were obtained at an ACE2 concentration of 4 μM and an RBD concentration of 8 μM. The sedimentation data were analyzed with SEDFIT software using the continuous sedimentation coefficient distribution model c(s) at a confidence interval of 0.95. The partial specific volume of protein, the buffer viscosity, and the density were calculated using SEDNTERP.

### Isothermal titration calorimetry

ITC experiments were carried out using MicroCal PEAQ-ITC (Malvern) at 25°C in a pH 7.0 buffer containing 50 mM Sodium Phosphate and 20 mM NaCl. RBDs at concentrations of 210 - 230 μM were titrated as 31 injections of 1.25 μl each into 30 μM of ACE2. The duration of each injection was 2.5 s and the interval between injections was 150 s. The data were analyzed using MicroCal PEAQ-ITC analysis software. For temperature dependence of interaction, RBDs were titrated into ACE2 incubated at different temperatures ranging from 10-30 °C with an interval of 5°C.

## RESULTS

### Sequence and structure comparison of the two RBDs

The structure and sequence comparison of the two RBDs is shown in Figure 1. The RBDs are composed of about 220 amino acids and show a sequence identity of about 70%. The ACE2 binding region is part of the receptor binding motif (RBM), which is towards the C-terminal region of RBD. RBM shows more sequence divergence and has a sequence identity of only about 50% between the two RBDs. In their crystal structures (0.42 Å RMSD determined using PyMol), RBM can be seen as an extended loop, with less regular secondary structures. Most of the ACE2 binding residues lie in the RBM and are shown in red font in Figure 1B. Twenty-three amino acids from both RBD1 and RBD2 make contact with the ACE2 protein. For RBD1, the interaction is more polar with 15 out of 23 residues being polar. For RBD2, the interaction involves more non-polar residues with 13 out of 23 residues being polar.

**Figure 1:**
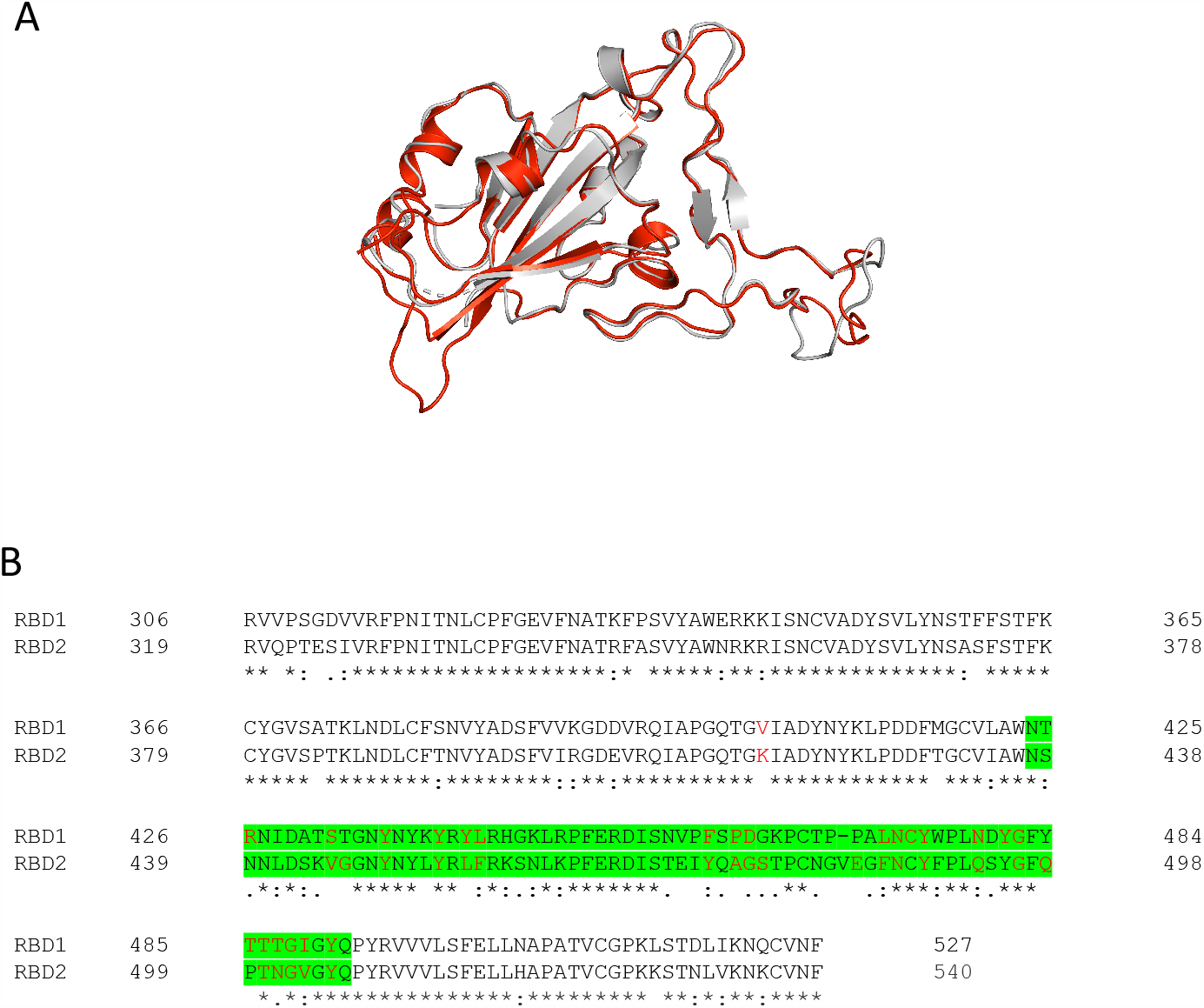
Sequence and structure comparison of RBD1 and RBD2. (A) A model of RBD1 and RBD2 overlap (obtained from PDB IDs 2AJF and 6M0J respectively). RBD1 is shown in black and RBD2 is shown in red. (B) The sequence alignment of RBD1 and RBD2 showing amino acid residues that forms the interface with ACE2. Identical residues are shown by asterisk (*), highly conserved residues are shown by colon (:), residues with low conversation are shown with periods (.) and dissimilar residues and gaps are represented as blanks. The RBM is highlighted green, and the interface residues are shown in red color.

### SARS-CoV-2 RBD shows lesser expression than SARS-CoV RBD

The proteins used in this study, ACE2 and receptor binding domains of SARS-CoV and SARS-CoV-2, were expressed in Expi293F cells. The differences in the level of expression between RBD1 and RBD2 were compared 3 days post-transfection from the culture supernatant using SDS-PAGE (Figure 2A). The expression was about 5 times lesser for RBD2 compared to RBD1, as calculated from Image-J analysis of the expression bands on SDS-PAGE. Both RBDs and ACE2 were recovered from the culture supernatant 5 days post-transfection and purified to homogeneity (Figure 2B).

**Figure 2:**
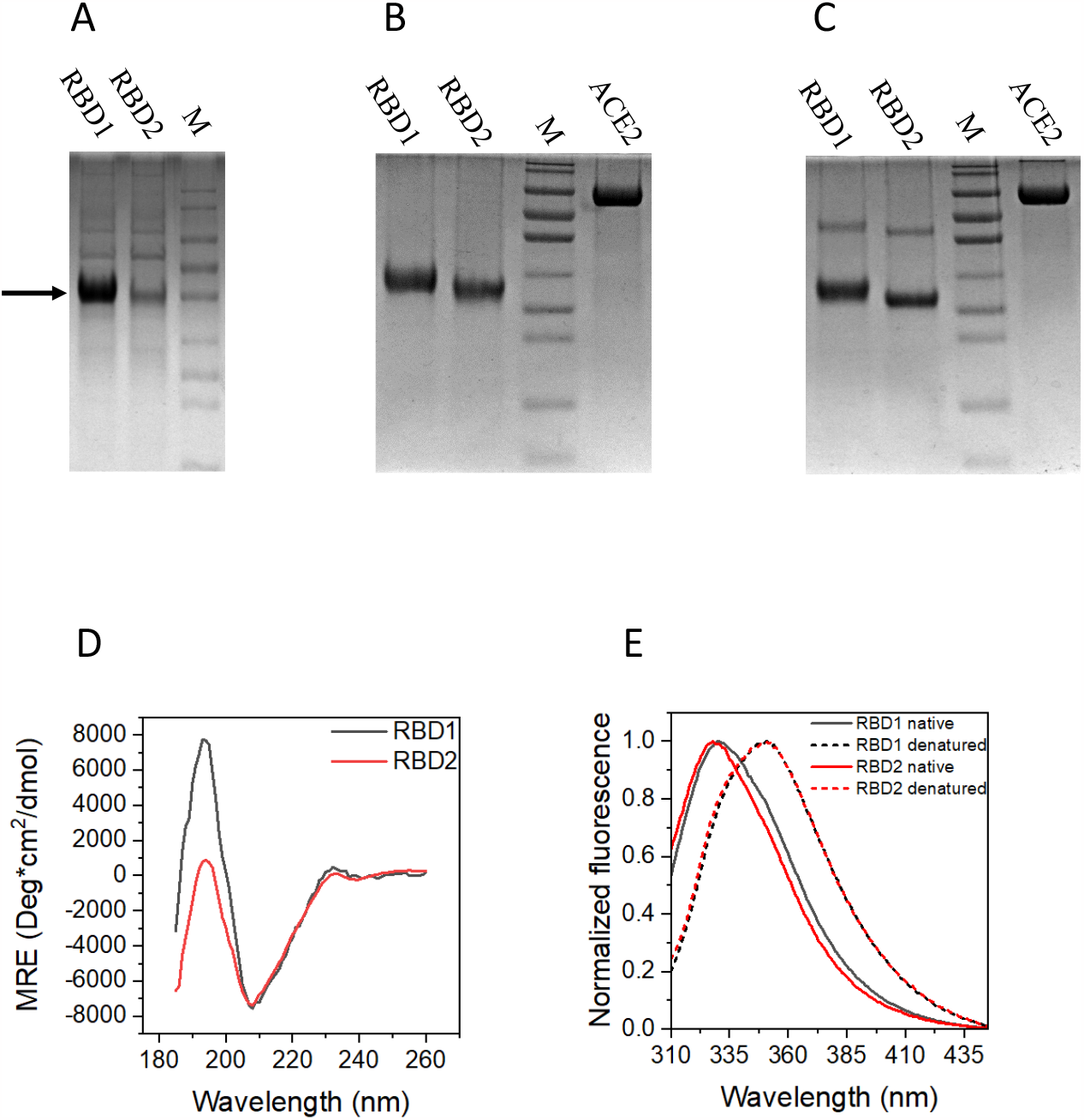
Protein expression, purification and characterization. (A) SDS-PAGE of the Expi-293 cell culture supernatant showing expression of RBD1 and RBD2, 3 days after transfection. The target protein band corresponding to RBDs is shown with an arrow. The purified RBD1, RBD2 and ACE2 in (B) reducing and (C) non-reducing conditions. Lane labeled M represents the molecular weight markers (180, 130, 100, 70, 55, 40, 35, 25, 15, and 10 kDa from top to bottom). (D) Far-UV CD spectra of RBD1 in black and RBD2 in red (E) Tryptophan fluorescence emission spectra of RBD1 in black and RBD2 in red. RBDs in native conditions are shown in solid line and in the denatured conditions (8.5 M urea) are shown in the dashed line.

Although the theoretical molecular weight of ACE2 is ∼70 kDa and that of RBDs is ∼25 kDa, the proteins run at a higher molecular weight due to glycosylation.(30-35) Analysis of the proteins on SDS-PAGE under non-reducing conditions reveals the presence of a small proportion of disulfide-linked dimer (Figure 2C). Both RBDs contain nine cysteine residues, with crystal structures showing the presence of 4 intramolecular disulfide bonds (PDB IDs: 6M0J for RBD2 and 2AJF for RBD1) leaving one cysteine free to form an intermolecular disulfide bond. The absence of higher oligomers other than dimer on the SDS-PAGE under nonreducing conditions (Fig. 2C) suggests the formation of correct disulfide bonds during intracellular folding with no disulfide scrambling during protein expression and purification.

### Purified SARS-CoV and SARS-CoV-2 RBDs have similar structures

CD and fluorescence spectroscopies were used to analyze the secondary and tertiary structures of RBDs. Far-UV CD spectra of RBDs show a prominent negative band at 208 nm and a positive band at 194 nm (Figure 2D). Deconvolution of CD spectra using BeStSel web software(36, 37) indicates the presence of a predominant β-sheet structure over α-helix structure with very similar secondary structural content for RBD1 and RBD2 (Table 1). The two RBDs differ in terms of the content of antiparallel beta-sheet and random coil structures, which accounts for the difference in the relative intensities of the positive CD band at 194 nm. The crystal structure as well as the structure prediction program show that about 45% of the sequence adopts random secondary structure and may be more dynamic or flexible than the structured regions of the two proteins. The ACE2 interaction is primarily mediated by the RBM, which shows more sequence divergence and also has an ill-defined secondary structure (Figure 1). This points towards the importance of flexible regions of RBDs in the ACE2 interaction. The sequence tolerance of well-defined secondary structure is usually low and only a few mutations are allowed without disrupting the fold of the protein. As dynamic regions of protein have more sequence tolerance, it can sample a wide range of mutations to fine-tune its function. Out of the 23 amino acid residues involved in ACE2 binding only eight residues are identical in RBDs, which further supports the hypothesis of high mutational frequency of less structured regions to fine-tune the function.

**Table 1.** Deconvolution of CD spectra of RBD1 and RBD2 using BeStSel web software.

| Secondary structure (%) | RBD1 | RBD2 |
| --- | --- | --- |
| Helix | 12.9 | 13.7 |
| Antiparallel | 29.9 | 25.0 |
| Parallel | 0.0 | 0.0 |
| Turn | 12.6 | 13.9 |
| Others | 44.6 | 47.3 |

Tryptophan fluorescence spectra provide information about the tertiary structure of proteins. RBD1 and RBD2 show strong fluorescence intensity having emission maxima around 330 and 328 nm, respectively (Figure 2E). Tryptophan emission maximum close to 320 nm indicates that the tryptophan residues are in the hydrophobic core of the proteins shielded from the solvent.

The fluorescence spectrum of RBD1 is slightly red-shifted compared to RBD, because of slightly different exposure of three tryptophan residues at 340, 423, and 476 amino acid positions in RBD1, compared to two tryptophan residues at 353 and 436 amino acid positions in RBD2. RBD1 tryptophan residues at positions 340 and 423 are conserved and appear at positions 353 and 436 for RBD2 (Figure 1B). The minor differences in the emission maxima could be because of the slightly different exposure of RBD1 tryptophan at the 476 position. In the RBD1 crystal structure, the extent of burial is highest for Trp340 followed by Trp423 and then Trp476, which is completely exposed to solvent. Solvent-exposed tryptophan residue shows red-shifted fluorescence maxima around 350 nm. Tryptophan fluorescence of RBD1 and RBD2 shows similar spectra (λ_max_ = 350 nm) after denaturation (8.5 M Urea).

### SARS-CoV-2 RBD is less stable than SARS-CoV RBD

Thermodynamic stability is an important factor that determines the protein structure, function, expression, and solubility.(38) We used thermal and chemical denaturation melts to assess the thermodynamic stability of RBDs. Figures 3A and 3B show the far-UV CD spectra of RBD1 and RBD2 with temperature. An increase in the CD signal of the negative band from 220-240 nm was observed with an increase in temperature with a larger change at 233 nm. In addition, a considerable decrease was observed in the intensity of the positive band at 195 nm with an isodichroic point at 212 nm, which is indicative of a structural transition between beta-sheet and random coil structures.(39) Thermal denaturation melts were plotted using ellipticity at 222 nm as a function of temperature (Figure 3C) and also using fluorescence emission at 320 nm with excitation at 280 nm (Figure 3D). The thermodynamic parameters calculated by the two-state unfolding model (Equation 1) are listed in Table 2. Fitting parameters indicate higher stability for RBD1 compared to RBD2. Midpoints of thermal denaturation T_m_ for RBD1 and RBD2 were found to be 65.6 ± 0.6 oC and 54.4 ± 0.3oC, respectively. Similar results were obtained for thermal denaturation of RBDs using fluorescence spectroscopy, which showed higher stability for RBD1 with a T_m_ value of 62.5 ± 0.6 oC compared to RBD2 with a T_m_ value of 59.3 ± 0.6 oC.

**Table 2.** Protein stability and aggregation parameters of RBDs obtained by fitting the data in Figures 3C-3G.

|  | **RBD1** | **RBD2** |
| --- | --- | --- |
| **Thermal denaturation** |  |  |
| $\Delta H_m$ (CD), kcal/mol | $58.7 \pm 4.3$ | $105.4 \pm 9.4$ |
| $T_m$ (CD), °C | $64.9 \pm 0.5$ | $54.4 \pm 0.2$ |
| $\Delta H_m$ (Fluorescence), kcal/mol | $94.5 \pm 2.6$ | $116.6 \pm 18.8$ |
| $T_m$ (Fluorescence), °C | $62.4 \pm 0.1$ | $57.9 \pm 0.4$ |
| **Urea denaturation** |  |  |
| $\Delta G^{\circ}_{unf}$ (CD), kcal/mol | $11.8 \pm 2.6$ | $3.0 \pm 0.5$ |
| m- value (CD), kcal/mol/M [urea] | $-1.8 \pm 0.4$ | $-0.6 \pm 0.1$ |
| $\Delta G^{\circ}_{unf}$ (Fluorescence), kcal/mol | $9.3 \pm 0.1$ | $7.6 \pm 0.2$ |
| m- value (Fluorescence), kcal/mol/M [urea] | $-1.5 \pm 0.0$ | $-1.3 \pm 0.0$ |
| **Aggregation kinetics** |  |  |
| Percentage amplitude (fast phase) | $23.3 \pm 0.3$ | $19.0 \pm 0.3$ |
| Time constant (fast phase), s | $364.1 \pm 7.4$ | $471.6 \pm 12.6$ |
| Percentage amplitude (slow phase) | $76.7 \pm 3.1$ | $81.0 \pm 0.4$ |
| Time constant (slow phase), s | $51,245 \pm 2,659$ | $17,950 \pm 195$ |

**Figure 3:**
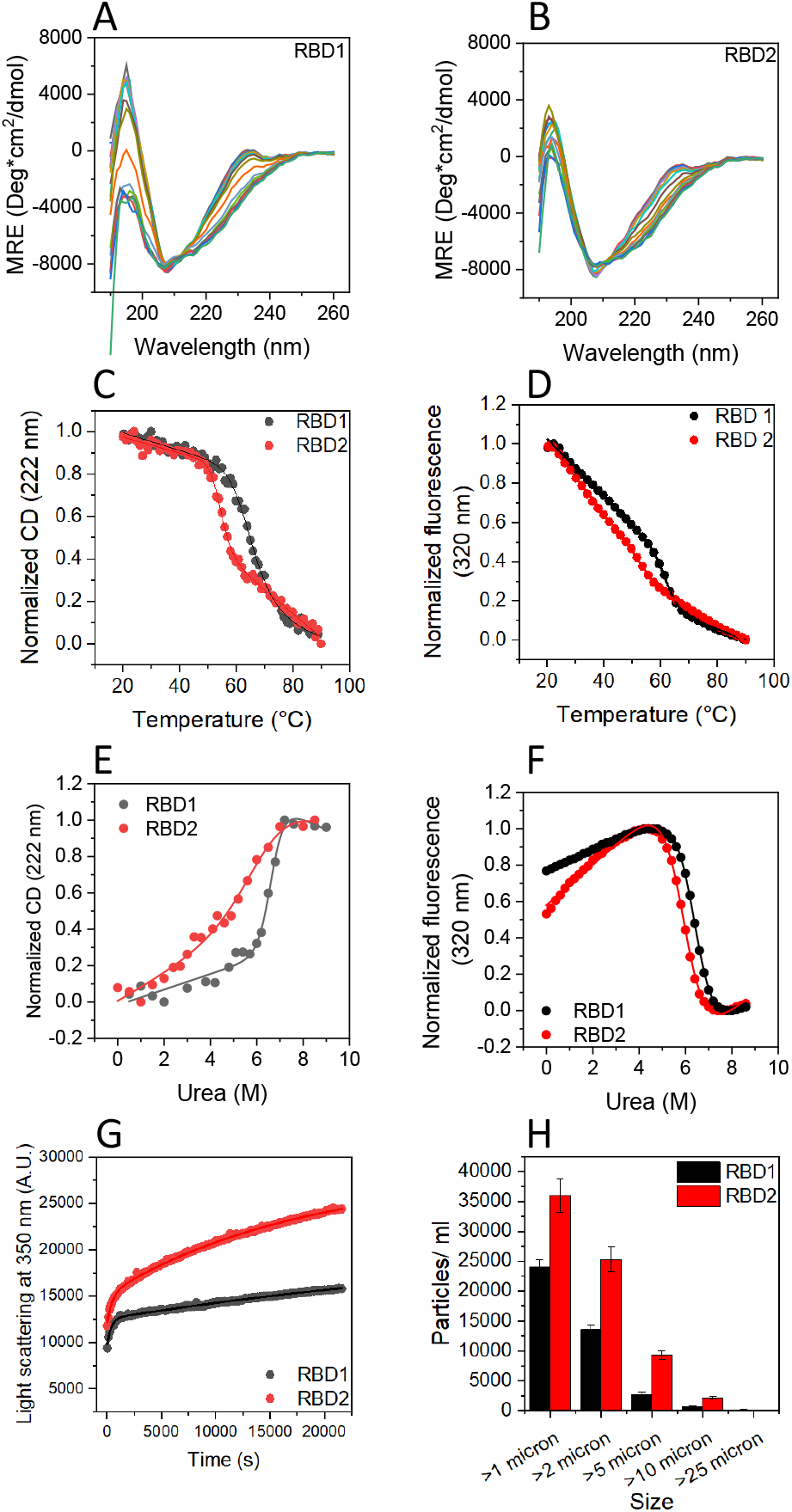
Stability and aggregation comparison of RBDs using denaturation melts and aggregation kinetics. Far-UV CD spectra of (A) RBD1 and (B) RBD2 at different temperatures ranging from 20 °C to 95 °C with 5 °C interval. Thermal denaturation melts of RBD1 and RBD2 monitored using (C) change in CD signal at 222 nm and (D) change in fluorescence emission at 320 nm with respect to temperature. Chemical denaturation melts of RBD1 and RBD2 monitored using (E) change in CD signal at 222 nm with respect to urea concentration and (F) change in fluorescence emission at 320 nm. For Figures 3C-3F, black and red represent the data corresponding to the RBD1and RBD2, respectively. (G) Aggregation kinetics of RBD1 and RBD2 monitored at 60 °C. (H) Size distribution of RBD1 and RBD2 particles counted using FlowCam after incubation at 60 °C for 30 h.

Urea denaturation profiles were obtained with both CD and fluorescence spectroscopy (Figures 3E and 3F). Both denaturation profiles showed higher stability for RBD1 compared to RBD2, consistent with the thermal denaturation melts. Because of increased absorbance of urea at high urea concentrations, CD signal at wavelengths lower than 220 nm could not be satisfactorily obtained. CD denaturation data at 222 nm could be fitted to a two-state unfolding model (Equation 3) with ΔG°_unf_ values of 11.8 ± 0.2 and 3.0 ± 0.6 kcal/mol for RBD1 and RBD2 respectively. Similarly, the fluorescence denaturation melts could be fitted well to a two-state unfolding model with ΔG°_unf_ values of 9.3 ± 0.1 and 7.6 ± 0.2 kcal/mol for RBD1 and RBD2, respectively.

### SARS-CoV-2 RBD has a higher aggregation propensity than that of SARS-CoV

The aggregation potential of the two RBDs was measured with an isothermal incubation at 60oC. Aggregation kinetics of both proteins could be fitted to a double-exponential equation with a fast phase and a slow phase. The fast phase was comparable for both proteins with a time constant of 364.1 ± 7.4 s and 471.6 ± 12.6 s for RBD1 and RBD2, respectively. The slower phase showed differences between the two RBDs with time constants of 51,245 ± 2,659 s for RBD1 and 17,950 ± 195 s for RBD2, indicating faster aggregation, forming larger aggregates for RBD2 compared to RBD1 (Figure 3G). Further confirmation of a greater extent of protein aggregation for RBD2 compared to RBD1 came from increased protein particles for RBD2 measured using FlowCam (Figure 3H). The data show a greater number of particles formed for RBD2 than for RBD1 in every size range. Both aggregation kinetics and the number of protein aggregate particles suggest a higher aggregation propensity for RBD2 compared to RBD1, consistent with the lesser stability of RBD2.

### RBD2 interacts strongly with ACE2 compared to RBD1

Both RBD1 and RBD2 interact with ACE2 through its RBM with a similar mode of interaction (Figures 4A and 4B). ITC was used to determine the strength of binding interactions between RBDs and ACE2. Thermograms indicate exothermic interaction between RBDs and ACE2, and the integrated heat changes fitted well to the one-site binding model. Fitted parameters indicate that RBD1 binds to ACE2 with a K_d_ of 64.8 ± 4.4 nM compared to that of RBD2 (K_d_ = 26.0 ± 3.0 nM), indicating that RBD2 binds strongly (∼2.5 fold change in K_d_) with ACE2 than RBD1. Another important difference between RBD1 and RBD2 towards ACE2 binding was the stichometry of interaction. The n-value for RBD1-ACE2 interaction was 0.6 ± 0.0 and that for RBD2-ACE2 interaction was close to 1 (n= 0.9 ± 0.0).

**Figure 4.**
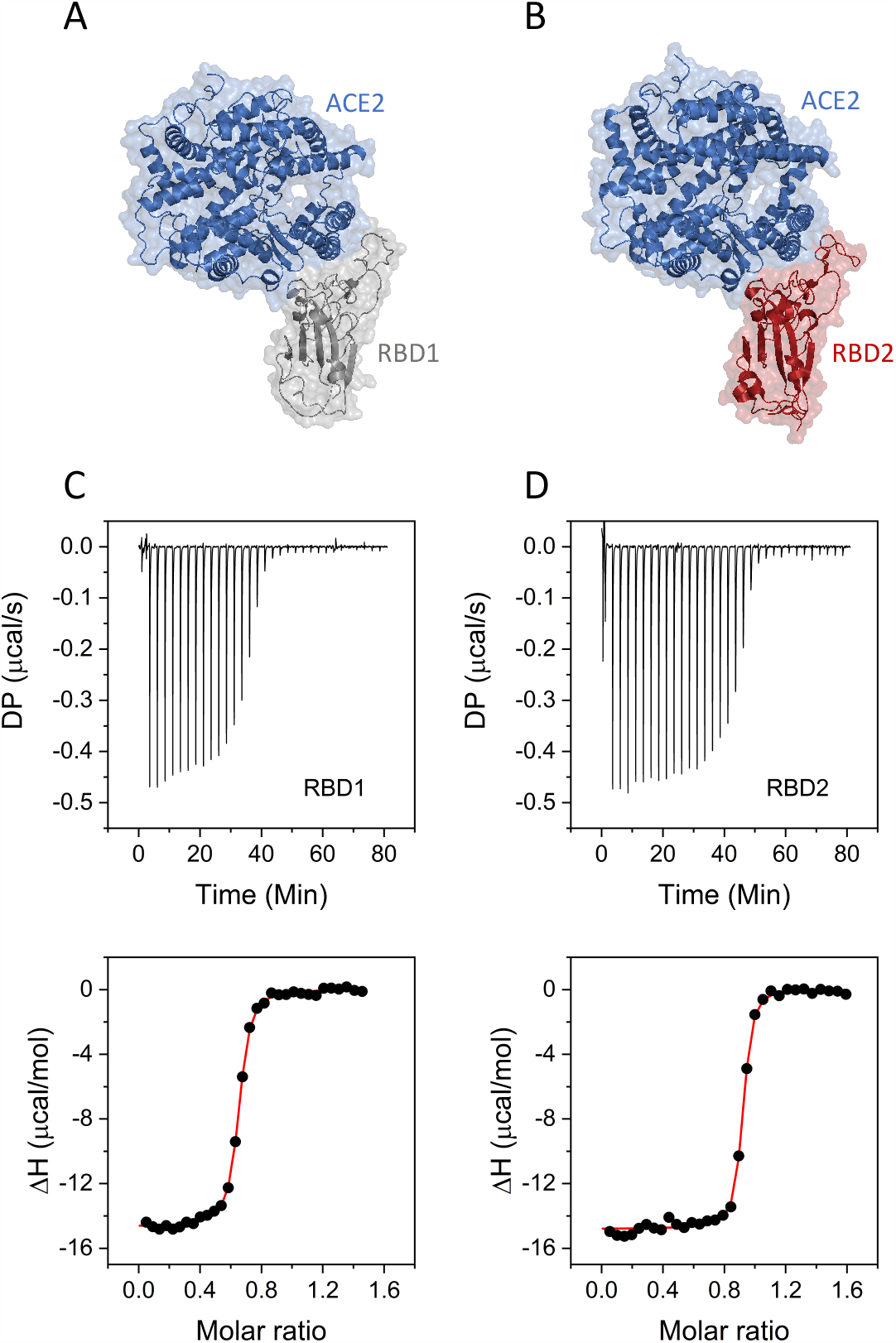
RBD-ACE2 interaction. A model of interaction of ACE2 with (A) RBD1; PDB ID 2AJF and (B) RBD2; PDB ID 6M0J. ACE2 is shown in blue, RBD1 is shown in black and RBD2 is shown in red. The ITC thermograms for interaction of ACE2 with (C) RBD1 and (D) RBD2 at 25 °C. The top panel shows the heat evolved with each injection. The bottom panel shows the integrated heat at each injection (black circles). The integrated heat plot was fitted to a single-site binding model to obtain the fit curve (red line) passing through the data points.

To determine the nature of ACE2 interactions between the two RBDs, the temperature dependence of binding affinity was further investigated. ITC thermograms recorded at five different temperatures ranging from 10oC to 30oC at 5oC intervals were shown in Figures 5A and 5B. The data suggests strong dependence of binding enthalpy (ΔH) on temperature, as evident from the ITC thermograms at five different temperatures. The thermodynamic parameters obtained from these ITC thermograms are shown in Table 3 and also shown in Figures 5C and 5D. The bar graph suggests a favorable increase in enthalpic contribution to binding with increasing temperature, which is accompanied by a proportional increase in unfavorable entropic component. This enthalpic-entropic compensation leads to a fairly constant ΔG over the temperature range studied. The slope of the linear dependence of ΔH with temperature was used to estimate the heat capacity ΔC_p_ values, which is an estimate of the accessible surface area (ASA) buried upon binding.(40) A comparison of the temperature dependence of ΔH along with the ΔC_p_ values for RBD-ACE2 interaction is shown in Figure 5E. The higher ΔC_p_ value for RBD1-ACE2 interaction suggests a larger surface getting buried upon interaction compared to RBD2-ACE2 interaction.

**Table 3.** Thermodynamic parameters for RBDs interaction with ACE2 at different temperatures obtained by fitting the data in Figures 5A and 5B.

|  |  | Temperature (°C) |  |  |  |  |
| --- | --- | --- | --- | --- | --- | --- |
|  |  | 10 | 15 | 20 | 25 | 30 |
| RBD1 | $\Delta H$ (kcal/mol) | $-8.8 \pm 0.1$ | $-10.2 \pm 0.1$ | $-12.5 \pm 0.1$ | $-14.6 \pm 0.1$ | $-16.8 \pm 0.1$ |
| | $K_d$ (nM) | $26.7 \pm 3.1$ | $25.8 \pm 3.6$ | $35.3 \pm 6.0$ | $64.8 \pm 4.4$ | $97.7 \pm 9.1$ |
| | N | $0.6 \pm 0.0$ | $0.6 \pm 0.0$ | $0.6 \pm 0.0$ | $0.6 \pm 0.0$ | $0.6 \pm 0.0$ |
| | $\Delta G$ (kcal/mol) | $-9.8 \pm 0.1$ | $-10.0 \pm 0.1$ | $-10.0 \pm 0.1$ | $-9.8 \pm 0.0$ | $-9.7 \pm 0.1$ |
| | $-T\Delta S$ (kcal/mol) | $-1.0 \pm 0.1$ | $0.2 \pm 0.1$ | $2.5 \pm 0.2$ | $4.8 \pm 0.1$ | $7.1 \pm 0.2$ |
| | $\Delta C_p$ (cal/mol/K) | $408.0 \pm 17.6$ | | | | |
| RBD2 | $\Delta H$ (kcal/mol) | $-9.9 \pm 0.1$ | $-11.5 \pm 0.1$ | $-12.7 \pm 0.1$ | $-14.8 \pm 0.1$ | $-15.7 \pm 0.1$ |
| | $K_d$ (nM) | $25.5 \pm 3.8$ | $17.4 \pm 3.6$ | $15.7 \pm 2.3$ | $26 \pm 3.0$ | $35.1 \pm 3.5$ |
| | N | $0.9 \pm 0.0$ | $0.9 \pm 0.0$ | $0.9 \pm 0.0$ | $0.9 \pm 0.0$ | $0.9 \pm 0.0$ |
| | $\Delta G$ (kcal/mol) | $-9.8 \pm 0.1$ | $-10.2 \pm 0.1$ | $-10.5 \pm 0.1$ | $-10.4 \pm 0.1$ | $-10.3 \pm 0.1$ |
| | $-T\Delta S$ (kcal/mol) | $0.1 \pm 0.1$ | $1.3 \pm 0.2$ | $2.2 \pm 0.1$ | $4.4 \pm 0.1$ | $5.3 \pm 0.1$ |
| | $\Delta C_p$ (cal/mol/K) | $297.6 \pm 18.1$ | | | | |

**Figure 5.**
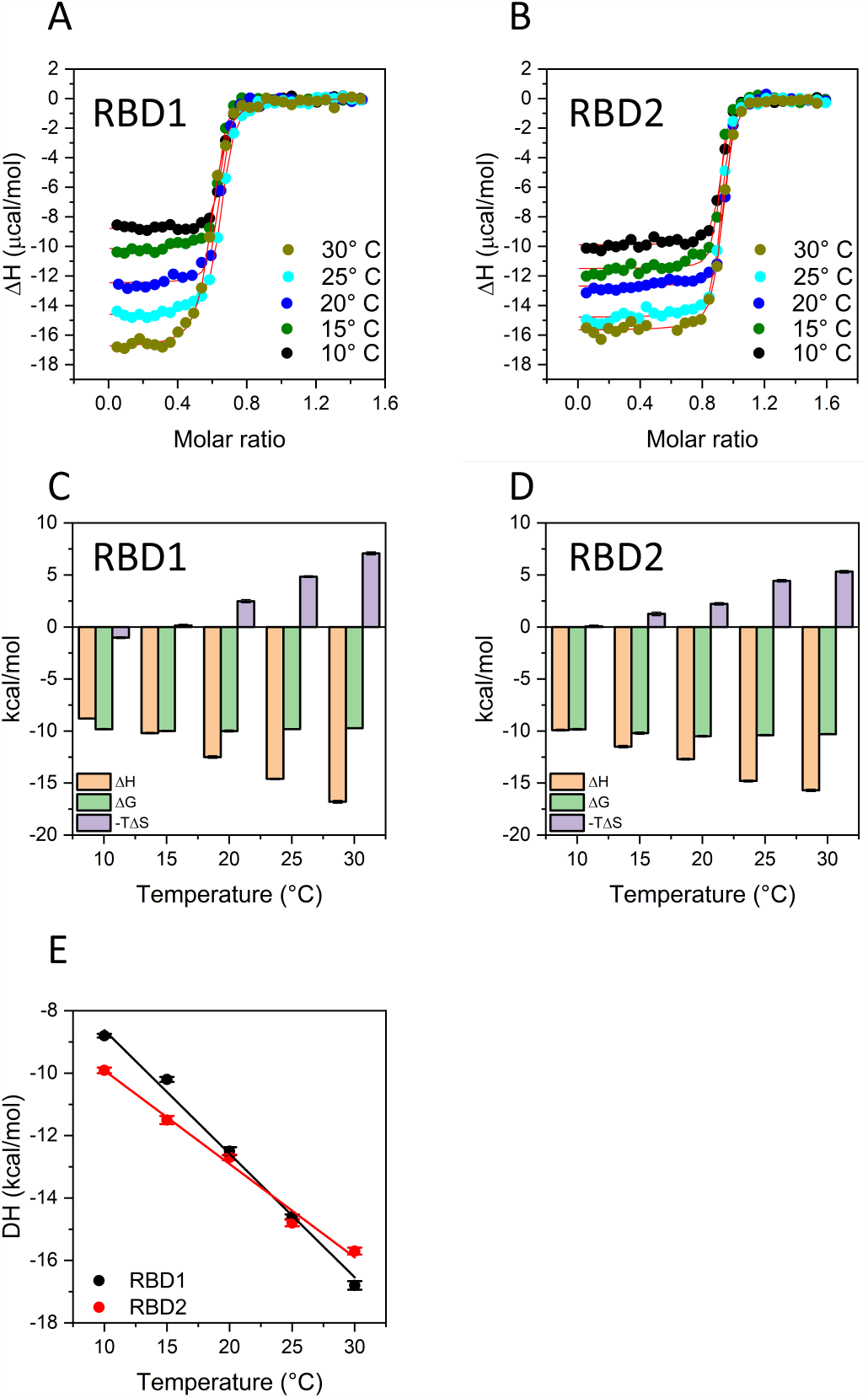
Temperature dependence of RBD-ACE2 interaction. The integrated heat plots are shown for temperature dependence of ACE2 interaction with (A) RBD1 and (B) RBD2. The thermodynamic parameters at different temperatures for ACE2 interaction with (C) RBD1 and (D) RBD2. (E) shows the comparison of linear temperature dependence of ΔH for RBD1 and RBD2 interaction with ACE2, the slope of which gives the ΔC_p_ values.

### Different oligomerization potential of RBD1 and RBD2 result in differential interactions of RBDs with ACE2

The SEC profiles were recorded for the ACE2 and RBDs individually and in their complex form (Figure 6). The SEC profiles of RBD1 and RBD2 at different concentrations are shown in Figures 6A and 6B respectively. Both proteins elute at an elution volume of around 10 ml. An important difference in the elution profile of the two proteins is the concentration-dependent shift of the elution peak maxima to lower volumes with increasing concentrations for RBD1, while no such change was observed for RBD2 (Figures 6A and 6B inset). The change in elution peak position with concentration is a hallmark of rapid equilibrium between species with different oligomeric status.(41, 42) Also, it is important to note that the width of distributions differs between RBD1 and RBD2. RBD1 eluted as a single peak, centered at an elution volume of 10.3 ml, with a very broad distribution indicating the presence of heterogeneity in the eluted protein sample. RBD2 on the other hand eluted as two distinct narrow peaks centered at 9 ml and 9.8 ml.

**Figure 6:**
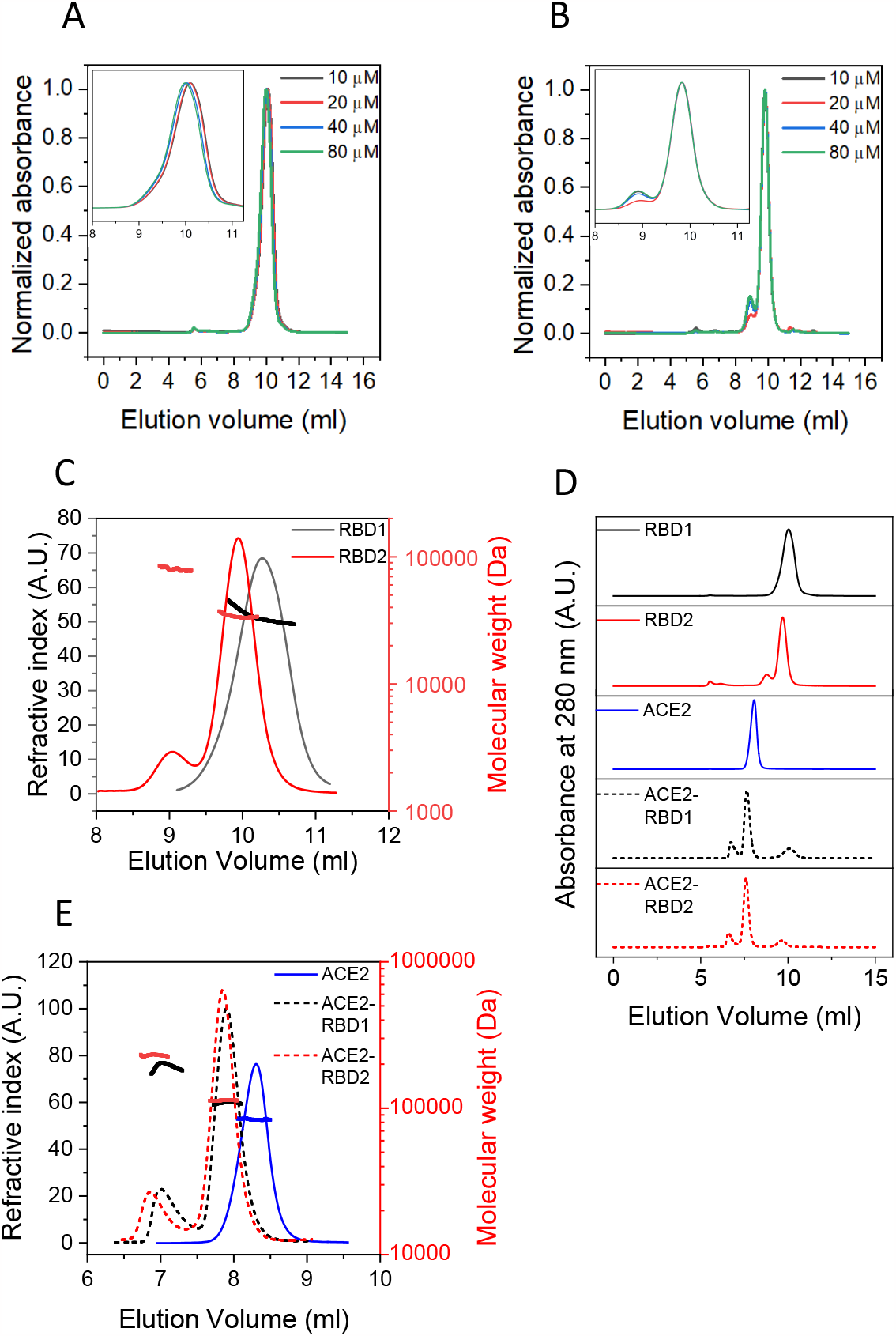
SEC-MALS analysis of interaction of RBDs with ACE2. The SEC profiles of (A) RBD1 and (B) RBD2 at different protein concentrations. The inset shows the SEC profile for the elution peak in zoomed-in version. (C) Comparison of the SEC-MALS profile of RBD1 and RBD2. Protein elution is followed as change in refractive index (vertical left axis) and plotted against elution volume and molecular weight (vertical right axis). The Horizontal curved lines on the graphs represents the calculated molecular weight. (D) Stacking graph of the SEC profile of RBD1, RBD2, ACE2, ACE2-RBD1 complex, and ACE2-RBD2 complex. (E) Comparison of the SEC-MALS profile of RBD1 and RBD2 after interacting with ACE2. For figures 5C-5E, ACE2 is shown in blue, while RBD1 and RBD2 are shown in black and red, respectively. The data for the complexes are shown in dashed lines. The molecular weight representations on the graphs follow the same color scheme.

The comparison of molecular weight estimates of RBDs from the SEC-MALS profiles is shown in Figure 6C. An important observation during the comparison of elution volumes is that RBD1 elutes later than RBD2, which points to either of the two possibilities: 1) The molecular weight of RBD1 is lower than RBD2 or 2) RBD1 is in a more compact conformation than that of RBD2. The molecular weights of the two proteins are not expected to be very different and if any, RBD1 appears to be of slightly higher molecular weight than RBD2 according to the SDS-PAGE analysis (Figures 2B and 2C). This strengthens the other possibility of RBD1 in a more compact conformation than RBD2, which is also supported by the unfolding m-values obtained from the urea denaturation melts for the two proteins (Figures 3E and 3F, Table 2). Consistent with the SEC profiles, the molecular weight calculations show that the distribution of molecular weight for the single SEC peak of RBD1 shows the presence of two species with molecular weights ranging approximately from 30 kDa to 45 kDa, which corresponds to the presence of monomers and dimers in the eluting protein that are not well resolved by the SEC column. The two peaks of RBD2, on the other hand, show two distinct molecular weight distributions. A low molecular weight species (average molecular weight 34.2 kDa) corresponding to RBD2 monomer and a high molecular weight species corresponding to RBD2 dimer (average molecular weight 80.5 kDa).

Compared to the two RBDs, ACE2 protein eluted as a narrow peak centered around an elution volume of 8.2 ml (Figure 6D). With SEC-MALS analysis (Figure 6E), ACE2 shows a single molecular weight distribution with an average molecular weight of ∼84.2 kDa, which indicates that ACE2 is a monomer in solution.

Analysis of the ACE2-RBD complexes by SEC was done at saturating concentrations of RBDs so that a lesser amount of free ACE2 is present in the solution, since free ACE2 and ACE2 bound to RBDs elute at similar elution volumes (Fig. 6D). The SEC profiles of both RBDs complexed to ACE2 revealed the presence of three main species: free RBDs, a low molecular weight ACE2-RBD complex and a high molecular weight ACE2-RBD complex (Figure 6D). Figure 6E shows the molecular weight distributions of ACE2 alone and in complex with RBDs. In complex with RBD1 and RBD2, ACE2 shows two distinct peaks - a major low molecular weight complex eluting at ∼7.8 ml and a minor high molecular weight complex eluting at ∼ 6.8-7.0 ml. The molecular weight distribution of the low molecular weight complexes with both RBD1 and RBD2 is fairly constant with average molecular weights of ∼108.8 and ∼113.3 kDa respectively. This means a 1:1 complex of ACE2-RBD is the major complex formed during the ACE2/RBD interaction. The molecular weight distributions of the high molecular weight complexes show differences among RBD1 and RBD2. RBD1-ACE2 complex shows a concentration-dependent change in its molecular weight. The molecular weight calculated at the peak apex is highest where the protein concentration is also high. As the protein concentration decreases towards both edges of the peak, the calculated molecular weight also decreases, indicating a rapid equilibrium between eluting species. The calculated molecular weight ranges from 172 - 205 kDa, suggesting a complex between 2 molecules of ACE2 and 1 or 2 molecules of RBD1. The origin of this dynamic complex could be attributed to the rapid monomer-dimer equilibrium seen for RBD1 (Figure 6A). The high molecular weight complex formed with RBD2, on the other hand, shows constant molecular weight distribution across the peak, with an average molecular weight of 230 kDa, which suggests a 2:2 complex between ACE2 and RBD2.

Similar to SEC-MALS experiments, the sedimentation coefficient of ACE2 and RBDs were calculated alone as well as in complex form using sedimentation velocity-AUC (SV-AUC) experiments (Figure 7A). Comparison of sedimentation coefficient distributions of RBDs (Figure 7B) shows two distinct peaks centered around 2.4 S and 3.8 S. The molecular weights of the two species calculated from SEDFIT analysis correspond to RBD monomers (33 kDa for RBD1 and 34 kDa for RBD2) and RBD dimers (62 kDa for RBD1 and 63 kDa for RBD2), respectively.

**Figure 7:**
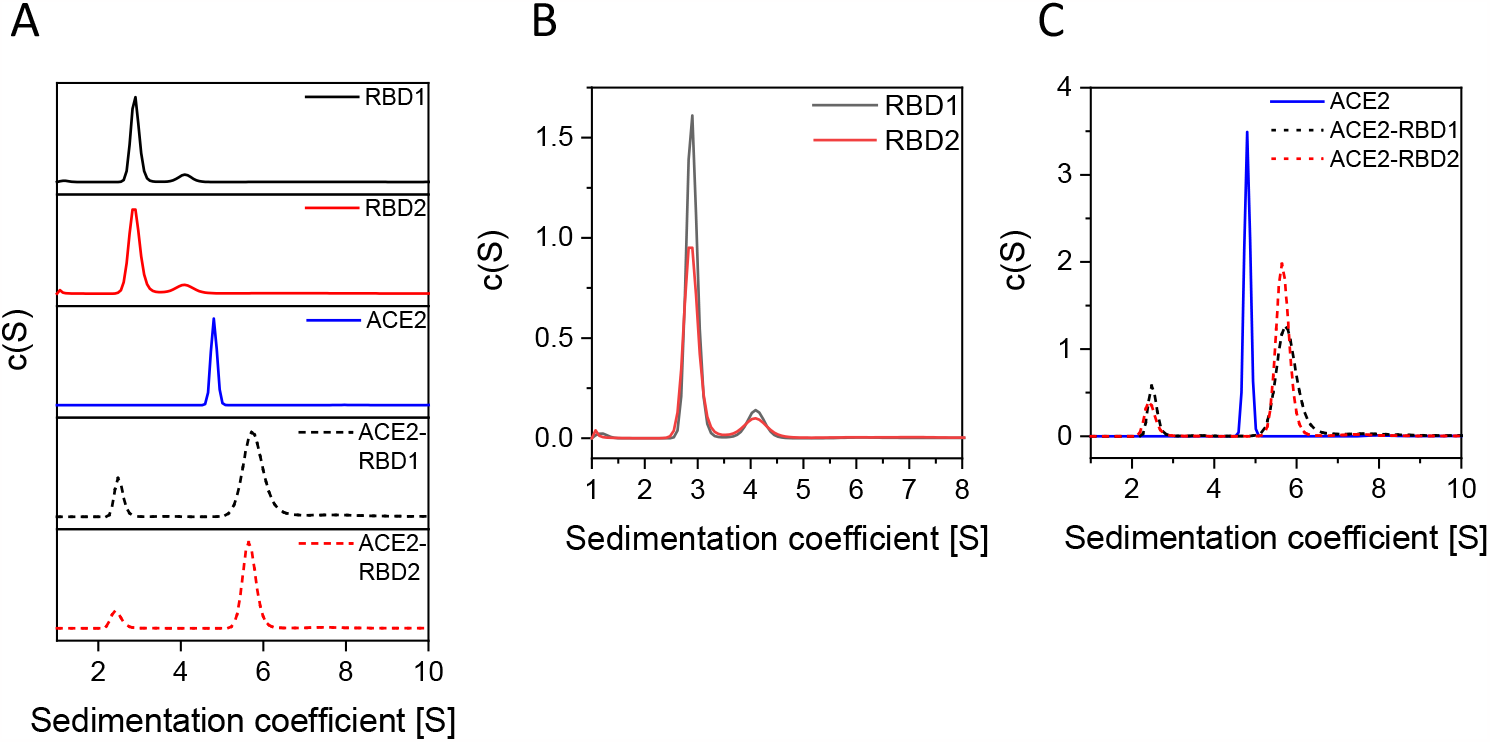
SV-AUC analysis of interaction of RBDs with ACE2. (A) Stacking graph of the SV-AUC profiles of RBD1, RBD2, ACE2, ACE2-RBD1 complex, and ACE2-RBD2 complex. (B) Comparison of the SV-AUC profiles of RBD1 and RBD2. (C) Comparison of the SV-AUC profiles of ACE2-RBD1 complex and ACE2-RBD2 complex with that of ACE2. ACE2 is shown in blue, while RBD1 and RBD2 are shown in black and red, respectively. The data for the complexes are shown in dashed lines

The sedimentation coefficient distributions of ACE2 and its complex with RBDs are shown in Figure 7C. Similar to SEC-MALS data, the sedimentation coefficient distribution of ACE2 shows a single narrow peak at 4.8 S with calculated molecular weight (using SEDFIT) corresponding to a monomer (83 kDa). Upon interaction with RBDs, the sedimentation coefficient increases to a higher value (5.7 S) and the width of the distribution also increases. The increase in the width of the distribution suggests the presence of heterogeneity in the sample. Another important observation is the greater width of distribution for ACE2/RBD1 complexes as compared to ACE2/RBD2 complexes. This means greater heterogeneity for ACE2/RBD1 complexes as compared to ACE2/RBD2 complexes as also seen in SEC-MALS experiments. Since SEC-MALS experiments were done at a higher protein concentration than AUC, we could see distinct species in SEC corresponding to differing stoichiometries.

## DISCUSSION

The outbreak of multiple coronaviruses has threatened human life during the last two decades. The first instance was SARS-CoV infection in 2002 followed by MERS-CoV infection in 2012. These outbreaks were not as contagious as the current pandemic caused by SARS-CoV-2, and the count of infected people was limited to a few thousand.(43) SARS-CoV-2 on the other hand has infected more than 687 million people, and despite its low pathogenicity compared to SARS-CoV and MERS-CoV, SARS-CoV-2 has caused more than 6.8 million deaths (https://covid19.who.int). The genomic similarity of SARS-CoV-2 with SARS-CoV and MERS-CoV is about 79% and 50% respectively,(44) which shows divergence of SARS-CoV-2 from these viruses. The closest genomic similarity of SARS-CoV-2 is found with viruses isolated from Bats and Pangolins (Bat-CoV-RaTG13 and pangolin-CoV-2020) with a genomic identity of 96% and 90% respectively.(45, 46, 6). Different theories on the origin of SARS-CoV-2 exist, although it is widely considered that human coronaviruses are derived from the bat reservoir.(5-8) SARS-CoV and MERS-CoV infected human hosts through intermediate hosts like civets and camels.(47, 48) The bat coronavirus is unlikely to be able to infect the human host directly due to low-affinity interaction with human ACE2, (49) which supports the possibility of an intermediate host of SARS-CoV-2 before infecting humans. To be able to infect human hosts, the virus needs to acquire features necessary for infecting humans. Efficient binding to ACE2 by mutating RBD of spike protein has been an important feature acquired by SARS-CoV and SARS-CoV-2. This acquisition of mutations in RBD could have happened through natural selection in animal hosts that have ACE2 remarkably similar to human ACE2 followed by zoonotic transmission to humans. According to a few reports, pangolins could have acted as the intermediate host for SARS-CoV-2 supported by the high-affinity interaction of pangolin coronavirus (pangolin-CoV-2020) with human ACE2 (K_d_ = 42.1 ± 10.0 nM).(49) Six critical residues in RBD are known to determine efficient binding with ACE2. All six of these critical residues are common between SARS-CoV-2 and pangolin-CoV-2020, whereas only one residue is common between SARS-CoV and SARS-CoV-2 RBD.

The RBD sequences of SARS-CoV and SARS-CoV-2 are divergent (Figure 1B) and show only 73% sequence identity. The acquisition of a common function of binding to ACE2 by both SARS-CoV and SARS-CoV-2 may be a result of convergent evolution, where similar functional traits are acquired in different lineages independently by natural selection. Convergent evolution is thought to be a product of positive selection,(50) and in this case, efficient ACE2 binding could have a selective advantage for the viruses that acquire it. This also means that future acquisition of ACE2 binding traits by other viruses is likely, which makes understanding the RBD-ACE2 interaction important. To compare the RBD sequence and its binding interaction with ACE2, we have selected the RBD of SARS-CoV (RBD1) and SARS-CoV-2 (RBD2), two viruses known to infect human hosts through interaction with ACE2. We have purified RBDs after expressing them in Expi293F cells, which are human cells and thus very closely match the natural infection scenario. The levels of protein expression vary for the two RBD constructs.

RBD1 shows ∼5 times higher expression than that of RBD2. Since the mammalian expression vector construct containing RBD1 and RBD2 contains a signal sequence that allows the translated protein to be secreted out of the cells, the proteins undergo the endoplasmic reticulum quality control while traversing the secretory pathway. The lower levels of secreted RBD2 compared to RBD1 could be due to the presence of a higher proportion of protein in a non-native or misfolded state. Proteins with non-native structures are prone to be cleared by the ER quality control mechanisms. Protein stability also seems to impact the levels of secreted protein through regulation by ER quality control mechanisms.(51) More stable proteins have more proportion of the protein in the native state and less proportion in the misfolded state, which is amenable to be cleared by ER quality control.

Protein stability is an important factor that determines the overall protein yields, especially for secretory proteins. It has been well established that more stable proteins have a higher protein expression level.(52) Protein expression levels have a direct bearing on the virus assembly and consequently infectivity. A higher expression would mean more protein for virus assembly. Our results show enhanced expression of RBD1 compared to RBD2, which could have a bearing on the infectivity of the two viruses, although expression levels of the complete spike protein and not just RBD would eventually determine the total amount of protein available for virus assembly. For example, it has been shown that the D614G mutation in SARS-CoV-2 spike protein increases virus infectivity by increasing the amount of functional spike protein by resisting cleavage by proteases during protein expression.(53) Protein stability can also have an impact on protein function. Proteins are in general marginally stable, and increased stability with mutations can sometimes results in a fitness penalty in the form of loss of function when more dynamic regions control function.(54) In the case of RBD-ACE2 interactions, increased stability of RBMs could be inversely proportional to the ACE2 binding function of RBD owing to its decreased flexibility.(55)

Protein stability also has important implications on the vaccine design strategy especially for the protein subunit vaccines. RBD is used as a source of antigen in multiple prospective vaccines against COVID-19.(19) Stability of vaccine formulation against protein aggregation is one of the important considerations for a successful vaccine. Both RBD sequences contain aggregation hotspots as confirmed by aggregation prediction servers like Waltz,(56) Tango,(57) Pasta,(58) and Aggrescan.(59) Stability of proteins can have an impact on their aggregation potential.(60) As RBD2 is less stable than RBD1, the aggregation potential of RBD2 is expected to be higher. Monitoring protein aggregation during thermal denaturation showed an increase in turbidity monitored by fluorescence at 350 nm for both RBDs. The midpoint of aggregation temperature was found to be higher for RBD1 than RBD2. Further, the kinetics of aggregation at 60oC was faster and generated more particles for RBD2 than RBD1 (Figure 3G and 3H). This means the use of RBD2 as antigen in vaccines is more challenging and any strategies that increase the stability of the protein or prevent aggregation would greatly benefit the vaccine formulations. As some viral variants with RBD mutations like K417T have higher stability compared to wild-type RBD,(61) use of such high stability variants for vaccine formulations could help design effective and safe vaccines.(62) It has also been shown that the compensatory effect of mutations could diminish the deleterious effects of other mutations,(61) so a careful selection of mutations in RBD, that can target different variants and at the same time have higher stability could be a key consideration from vaccine design point of view.

Protein oligomerization plays a significant role in protein function and stability. Our results suggest that the two RBDs show differences in the oligomerization potential. SEC-MALS experiments showed the differences in the elution profile and molecular weight distributions for the two proteins. The elution peak was much broader for RBD1 suggesting heterogeneity, supported by molecular weight distributions determined from SEC-MALS analysis (Figure 5). The impact of this heterogenous distribution of the RBD1 native state is recognized in its function as well. ACE2-RBD1 complex as probed by SEC-MALS showed the presence of heterogenous high molecular weight species that amounted to a 1:1 complex and a dynamic complex with two molecules of ACE2 interacting with one or two molecules of RBD (Figure 5). AUC data was consistent with the SEC-MALS data and showed broader peak distribution for the ACE2-RBD1 complex (Figure 5). Consistent with these results, the ITC data also showed differences in the stoichiometry of interaction of the two RBDs with ACE2. The n-value for RBD2-ACE2 interaction was close to 1, but that for RBD1-ACE2 interaction was 0.6 suggesting sub-stoichiometric interaction of RBD1 with ACE2, which could be explained by a rapid monomer-dimer equilibrium of RBD1 in its native state. Although the thermal and chemical denaturation data could be fit to a two-state unfolding model, they may not truly represent two-state unfolding. The thermal denaturation melts were not completely reversible and thus can only be used as a qualitative measure of the T_m_ value and apparent stability of the proteins. The chemical denaturation melts fit well to a two-state unfolding model; however, the possibility of other folding mechanisms involving monomer-dimer equilibrium cannot be ruled out. In addition, the denaturation melts of both RBDs (Figure 3E) show a sloped baseline, which might suggest a non-2-state unfolding behavior.

## CONCLUSIONS

Since the emergence of COVID-19 pandemic, there has been a heightened focus on understanding what new features bat CoVs can acquire to transition into human CoVs.(63-66) Among the three SARS human CoVs that resulted in a large number of infections and increased mortality rate, SARS-CoV and SARS-CoV-2 follow the same ACE2 receptor pathway to enter human cells. In this study, we have compared the receptor binding domains of SARS-CoV and SARS-CoV-2 in terms of their structure, stability, aggregation, and function. The amino acid sequences of the two RBDs are very divergent and share little sequence similarity, especially in the RBM, which is the functionally relevant region of the protein. These sequences might have evolved from a common ancestor and could have undergone convergent evolution to acquire the common function of ACE2 binding, although through quite different evolutionary histories. The two RBDs are very similar in secondary and tertiary structure and adopt a similar beta-sheet-rich fold with a significant contribution from the random coils. The stability of the two proteins is significantly different with a 10oC difference in the T_m_ of the two proteins as measured using CD. When measured by urea denaturation, the thermodynamic stability also differs between the two RBDs with ΔΔG°_unf_ of about 2-10 kcal/mol depending on what spectroscopic probe was used (Table 2). The aggregation propensities of the two proteins are also different with less stable RBD2 prone to increased aggregation. The native states of the two proteins also show differences in their oligomerization, with RBD1 having more propensity to form dimers and show rapid monomer-dimer equilibrium, which also affected its interaction with ACE2 protein. The RBDs also showed differences in their stoichiometry of interaction with ACE2, with RBD1 showing more heterogenous distribution of ACE2-RBD complexes owing to a monomer-dimer equilibrium of RBD1. Finally, the ACE2 binding affinity of RBD2 is higher than that of RBD1, which can be owed to fine-tuning specific interactions in the ACE2-RBD complex by optimizing the sequence of RBM. In addition, these results highlight significant differences in the RBDs of SARS-CoV and SARS-CoV-2 with respect to structure, stability, aggregation, and function. These differences should be important in understanding the evolution of the RBD of coronaviruses in general and the structural basis of ACE2-RBD interactions that determines the spillover of different coronaviruses from animal sources and endanger humans. It will be interesting to see whether any other SARS-CoVs emerging in the future follow the same biophysical principles examined here for the two SARS-CoVs that infected humans. Knowing the properties of RBD could also have important implications on the vaccine design strategies for SARS-CoVs that rely on the use of RBD as an antigen in protein subunit vaccines.

## AUTHOR CONTRIBUTIONS

V.U., S.P., and K.M.G.M. designed research; V.U., S.P., A.L., and C.P. performed research; V.U., S.P., and K.M.G.M. analyzed data; V.U., S.P., and K.M.G.M. wrote the manuscript.

## ACKNOWLEDGMENTS

This work was supported by an Associate Dean of Research seed grant program from the University of Colorado Skaggs School of Pharmacy and Pharmaceutical Sciences.

